# Isolation and identification of AMF species from selected medicinal plants from BHU Campus

**DOI:** 10.64898/2026.04.20.719602

**Authors:** Subhesh Saurabh Jha, L.S. Songachan

## Abstract

The objective of this study was to investigate Arbuscular Mycorrhizal Fungi (AMF) associations in selected medicinal plants. In this study 15 commonly used medicinal plants viz., *Abutilon indicum* (L.) Sweet*, Centella asiatica* (L.) Urb*, Piper longum*(L.)*, Terminalia bellerica* (Gaertner) Roxb*, Tinospora cordifolia* (Wild.) Miers, *Withania somnifera, Azadirachta indica* A. Juss., *Asparagus racemosus* Willd., *Andrographis paniculata* (Burm. Fil.) Nees, *Ocimum sanctum* L. *Eclipta alba*, *Mentha arvensis*, *Elettaria cardamomum*, *Bacopa monnieri* and *Mimosa pudica* were investigated for AMF colonization in the form of arbuscules, vesicles and hyphae from their roots and rhizosphere soil. The rhizosphere soil and root of the commonly used medicinal plants were procured from Banaras Hindu University (BHU). From the study it was clear that AMF spores are abundantly available in the rhizosphere of the plants chosen for this study with spores of Acaulosporaceae and Glomeraceae family being dominant and *Funneliformis mossae* having the highest relative abundance and isolation frequency among all the AMF species.

## Introduction

Medicinal plants have been used for centuries across diverse cultures to treat a wide spectrum of ailments. These plants contain natural compounds that possess healing properties and offer relief from various conditions. While widely celebrated for their medicinal benefits, it is crucial to exercise caution when using these plants, particularly in large quantities or with concentrated forms such as in essential oils or extracts. For this study 15 commonly used medicinal plants were considered for studying the composition of mycorrhizal associations in their rhizosphere. A brief discussion about some of the medicinal properties of these plants are discussed below:-1.) Indian mallow (*Abutilon indicum*)*-* It is a small shrub in the family Malvaceae, native to tropical and subtropical regions and is used in traditional medicine for variety of ailments. 2.) Kalmegh (*Andrographis paniculata*)*-Andrographis paniculata* is a medicinal herb that is native to India and Sri Lanka. It is a member of the Acanthaceae family of plants used in traditional ayurvedic/Chinese medicine. 3.) Shatavari (*Asparagus racemosus*)*-* Shatavari belongs to the family of Asparagaceae, it is a climber having phylloclades. It is used to treat gastrointestinal issues and also in maintaining a healthy digestive tract enhancing body resistance to stress and disease, acting as an immunomodulator. Due to its anti-inflammatory properties, it is also used in treating respiratory ailments like bronchitis and chronic coughs. 4.) Neem *(Azadirachta indica*)*-Azadirachta indica* also known as the neem tree, is native to India and Southeast Asia. It is a member of the Meliaceae family. Neem is widely known for its antibacterial and antifungal properties, making it effective in treating various skin conditions. 5.) Brahmi (*Bacopa monnieri*)*-* Brahmi belongs to the family of Plantaginaceae. It has been used in ayurvedic medicines and is celebrated for its cognitive-enhancing properties. 6.) Gotu kola (*Centella asiatica*)*-*It is an herbaceous, perennial plant belonging to the family of Apiaceae. Gotu kola is best known for its ability to heal wounds, treat burns, and improve skin conditions. It promotes collagen production, which is crucial for wound healing and maintaining skin elasticity.7.) Bhringraj (*Eclipta alba*)*-* Belonging to the family of Asteraceae, it is believed to prevent hair loss, stimulate hair growth and combat premature greying. Bhringraj oil is used for hair treatment. *Eclipta alba* is also known for its hepatoprotective properties, and is often used in treating cirrhosis.8.) True cardamom *(Elettaria cardamomum*)*-* Belonging to the family of Zingiberaceae and cultivated widely in tropical regions and reportedly naturalized in Reunion, Indochina and Costa Rica. It has antioxidant and diuretic properties. It is well known for its ability to aid in digestion, reduce bloating, and treat intestinal spasms. It is often used in traditional medicine to combat nausea and vomiting. Due to its antibacterial properties and refreshing aroma, it is also often used in treating bad breath and preventing cavities. The spice is believed to be beneficial for those with asthma or other respiratory issues due to its anti-inflammatory properties. 9.) Wild mint (Mentha *arvensis*)*-Mentha arvensis* belongs to the family of Lamiaceae. It has a circumboreal distribution being native to the temperate regions of Europe and western and central Asia. It has been used for various purposes, owing to its menthol content and other beneficial compounds. One of the most common uses of wild mint is in aiding digestion. It has also been found to be effective in relieving symptoms of the common cold, such as congestion, cough, and sore throat.10.) *Touch-me-not (Mimosa pudica*)*-Mimosa pudica also known as the* touch-me-not plant is a creeping annual/perennial flowering plant belonging to the family of Fabaceae. The plant has been used traditionally for its wound healing properties. It is believed to promote cell regeneration and reduce inflammation, making it beneficial for treating cuts and wounds. In some traditional medicine systems, *Mimosa pudica* is used as an anti-venom agent, particularly for snake bites. The plant is used to treat gastrointestinal disorders like diarrhea and dysentery due to its antispasmodic and astringent properties.11.) Tulsi *(Ocimum sanctum*)*-* Tulsi is known as an adaptogen, meaning it helps the body adapt to stress and promotes mental balance. It is believed to strengthen the immune system and is often used to prevent and treat colds, coughs, and other respiratory ailments. It is also used to promote heart health, potentially lowering blood pressure and cholesterol levels. Tulsi can aid in digestion and help treat stomach problems like bloating, gas, and cramping. The plant has powerful antioxidant properties. It shows antimicrobial activity against a range of bacteria, viruses, and fungi. It may help in regulating blood sugar levels, making it beneficial for people with diabetes. It’s often used for its detoxifying properties, helping to purify the blood and remove toxins from the body.12.)Long pepper (*Piper longum)-* Long pepper, sometimes called Indian long pepper or *pippali*, is a flowering vine in the family Piperaceae. It is known for its ability to stimulate the appetite and enhance digestion. It is also commonly used in treating respiratory conditions like cough, bronchitis, and asthma due to its expectorant properties. One of the most significant properties of long pepper is its ability to increase the absorption and effectiveness of other drugs and herbs, making it a valuable component in various herbal formulations.13.) Baheda (*Terminalia bellerica*)*-T. bellerica is* a large deciduous tree in the Combretaceae family. It is highly valued in ayurvedic medicine for its diverse health benefits. It is commonly used for treating respiratory conditions like bronchitis, asthma, and sore throat due to its astringent and anti-inflammatory properties. It is beneficial in improving digestion and treating digestive disorders, including constipation, which is why it’s a key ingredient in triphala. 14.) Giloy *(Tinospora cordifolia)-* Giloy is renowned for its immune-boosting properties. It is believed to enhance the body’s resistance to infections and illnesses. It’s used to reduce inflammation, making it beneficial in treating conditions like arthritis and gout. It aids in digestion and is used in treating digestive disorders such as hyperacidity, colitis, and worm infestations. It is effective in reducing fever and is often used as a natural remedy for dengue and malaria fevers. 15.) Ashwagandha *(Withania somnifera)-* Ashwagandha, is an evergreen shrub in the Solanaceae or nightshade family that grows in Nepal, India, the Middle East, and parts of Africa. One of its most well-known uses is as an adaptogen, a substance that helps the body cope with stress. It’s been shown to reduce levels of stress and anxiety. Ashwagandha is believed to enhance memory and brain function, making it a popular supplement for cognitive health. The herb has anti-inflammatory and pain-relieving properties, which are useful in treating conditions like arthritis.

All these plants were investigated for AMF colonization. AMF are important for the health of many plant species and are found in a wide range of ecosystems, including grasslands, forests, and agricultural systems. They are particularly important in ecosystems with low levels of available nutrients, as they help plants to access these nutrients and improve their growth and survival. AMF are also believed to have a number of ecosystem-level effects, such as improving soil structure and promoting plant diversity. Effective utilization of AMF though requires a deep understanding of the complex interactions between plants, microbes, and soil. These multitrophic relationships demand interdisciplinary research using molecular, biochemical, and physiological approaches (Jha and Songachan, 2022). There are many different species of AMF, and their specific roles and interactions with plants can vary. However, in general, AMF are considered to be important components of healthy ecosystems and have the potential to play a role in sustainable growth and development of plants.

AMF associations are characterized by small, tree-like structures called arbuscules within root cells. The arbuscules are highly branched structures that are able to penetrate deep into the cell walls and cytoplasm of the root cells, allowing the fungus to absorb nutrients and water that it obtains from the soil, helping the plant to grow and thrive and vesicles which are specialized cells representing the dormant stage in AMF’s life-cycle and has the primary function of storage (Jha and Songachan 2020).

Plant diversity plays a crucial role in shaping AMF diversity. Studies have shown that diverse plant communities support a higher diversity of AMF (Mahanta *et al*., 2018). Different plant species exhibit varying degrees of specificity towards particular AMF species, and the presence of diverse plant species can provide a wider range of niches and resources for AMF, leading to increased fungal diversity (Wang *et al*.,2018).

The identification of AMF can be challenging due to their complex life cycle and the absence of easily observable morphological features. Isolation and culture techniques involve extracting AMF spores or mycelium from soil or plant roots and establishing pure cultures (Landis *et al*., 2004; Gai *et al*., 2006). These cultures can then be subjected to morphological and molecular analyses for identification. Traditional morphological methods involve examining the structures of AMF under a microscope. The key morphological features used for identification include spore size, shape, color, and wall ornamentation. Molecular methods, particularly DNA-based techniques, have revolutionized AMF identification by providing greater accuracy and resolving power. These methods involve the extraction of DNA from fungal samples, followed by amplification and sequencing of specific target regions, such as the small subunit (SSU) rRNA gene or the internal transcribed spacer (ITS) region. Phylogenetic analysis of the obtained sequences can then be used to identify the AMF species (Reddy *et al*., 2005).

## Methodology

### Study site and sampling

The study was conducted at BHU Campus, Varanasi. In this study 15 commonly used medicinal plants viz., *Abutilon indicum, Andrographis paniculata, Asparagus racemosus, Azadirachta indica, Bacopa monnieri, Centella asiatica, Eclipta alba, Elettaria cardamomum, Mentha arvensis, Mimosa pudica, Ocimum sanctum, Piper longum, Terminalia bellerica, Tinospora cordifolia and Withania somnifera* were investigated for AMF colonization in the form of arbuscules, vesicles and hyphae from their roots and their rhizosphere soil was isolated for AMF spores. The sample were collected in sterilized plastic bags and transported to the laboratory for further analysis. The study showed that AMF spores are abundantly available in the rhizosphere of the plants chosen for this study with spores of Acaulosporaceae and Glomeraceae family being dominant and *Funneliformis mosseae* having the highest relative abundance and isolation frequency among all the AMF species.

### Estimation of AMF colonization

Root colonization was quantified using the technique of microscopic examination of root samples. Plant roots were carefully collected, washed to remove soil particles, cut into 1cm segments. The root segments were then heated with 10% KOH for about 10 mins. at 90°C, after which it was washed thoroughly to remove all traces of chemicals. The roots were then stained with trypan blue to visualize the fungal structures. Microscopic observation was done to identify and measure the presence of AMF structures such as arbuscules, vesicles, and hyphae within the root tissue. The percentage of root length colonized by AMF was calculated based on these observations.

### AMF spore isolation, identification and enumeration

AMF spores were isolated by technique known as wet sieving and decanting method (Gerdemann and Nicolson, 1963).According to protocol, 100g of rhizosphere soil was taken and washed under running tap water through a series of sieves, the soil mixture was agitated vigorously to free the AMF spores from soil and allowed to settle for 15-45 minutes and the supernatant was decanted through standard sieves of 300µm,150µ m and 45µm.The small particles left in the second tier (150µm) were collected along with tap water and kept it in a beaker for 10-15mins without any disturbance to settle down the debris below and spores on the surface. The water sample was then passed through a filter paper and the spores present along with debris were trapped on the paper and the water sample was then discarded. The filter paper was then spread on the petri dish and spores were counted using stereo microscope. Sporocarps and spore clusters were taken as single unit. AMF spores were picked up using a needle and mounted in PVLG. AMF spores were identified based on their morphological characteristics such as size, shape, wall ornamentation, colour, presence or absence of subtending hyphae, etc. Spore density and species richness were expressed as number of AM fungal spores and numbers of AM fungal species in 100g soil sample.

### Soil Physio-chemical analysis

Soil moisture was determined by drying 10g fresh soil at 105°C for 24h in hot-air-oven. Soil pH was determined using a digital pH meter. Organic carbon was analysed by colorimetric method and available phosphorus by molybdenum blue method.

### AMF trap culture

AMF trap culture was done using the methods of INVAM. AMF inoculum (rhizosphere soil) was mixed in the ratio of 1:1(v/v) with autoclaved coarse sand (sieved through 30 mesh size BBS) and placed in pots. The pots were seeded with *Sorghum bicolor* as host plant. For each treatment 3 replicates in a pot were prepared. Pot of non-mycorrhizal control were also raised. In the non-mycorrhizal pot, sterilized soil+ sterilized sand (1:1 v/v) without AMF inoculum were kept with all other conditions same. The cultures were grown in a greenhouse at 20±5 with 60 % relative humidity. The pots were watered at regular intervals and were arranged on a greenhouse bench in a completely randomized design with three replicates each. Half-strength Hoagland’s nutrient solution (Hoagland and Arnon,1938) was provided to the plants at fortnightly interval. After four months of growth cycle, the pots were left to dry undisturbed with a fairly stable temperature so that drying period is not too rapid, after which, the dried shoots were cut at ground level (Figure 8).

**Figure 1.**
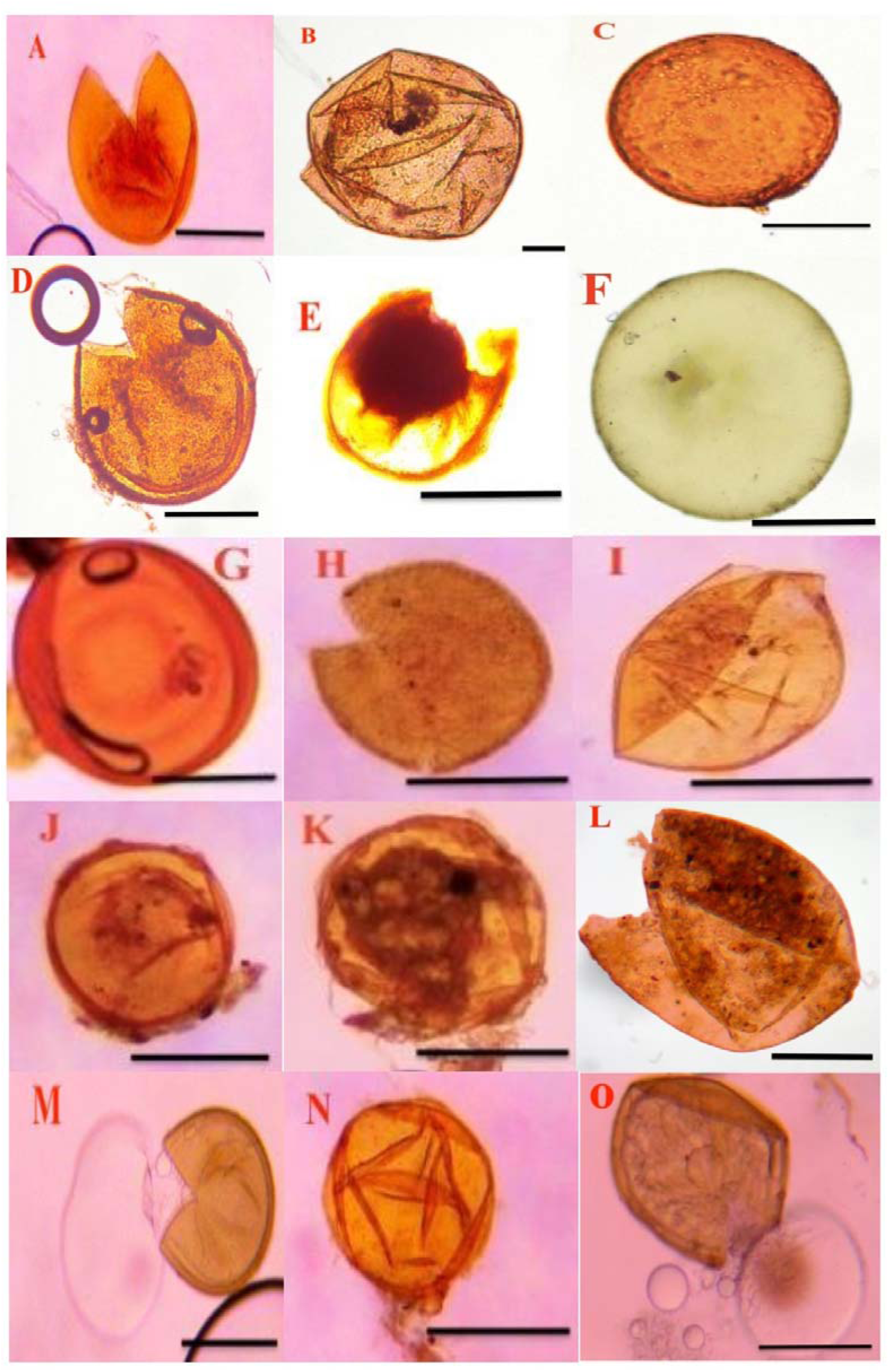
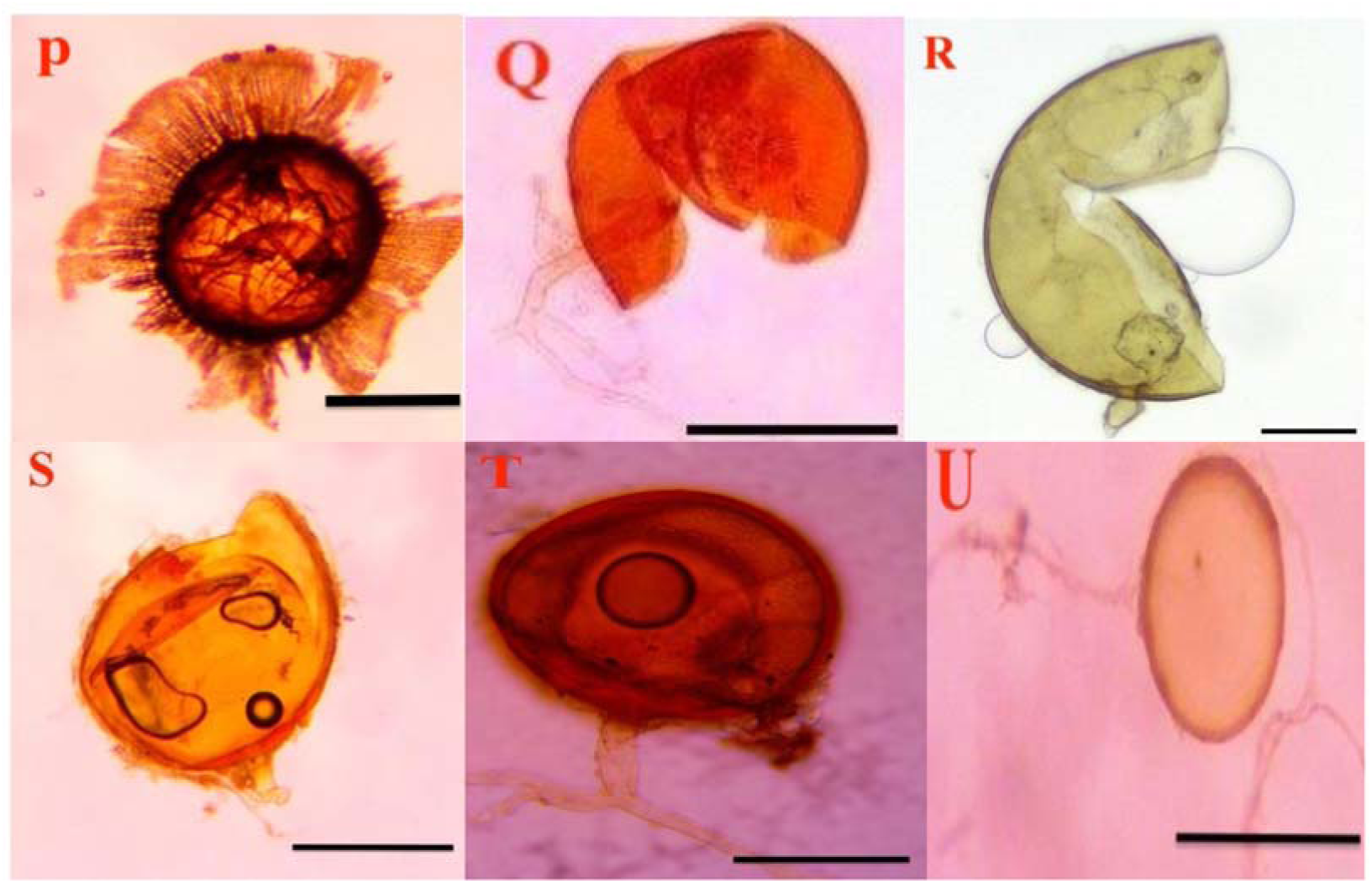
A-O-(Acaulosporasps.). A.)A.bireticulata, B.)A.scrobiculata, C.)A.spinosa, D.)A.koskeii, E.)A.lacunosa, F.)A.longula, G.)A.mellea, H.)Acaulospora sp.1, I.)Acaulospora sp. 2, J.)Acaulospora sp. 3, K.) Acaulospora sp. 4, L.)Acaulospora sp. 5, M.)A.saccate, N.)A.myriocarpa O.)A.delicata, P.)Ambisporaappendiculata, Q.)Funneliformismosseae,(R-U-Gigaspora sps.) R.) Gigaspora albida, S.)Gigaspora decipiens,T.)Gigaspora sp. 1, U.) Gigaspora margarita (Scale 40 µm).

**Figure 2.**
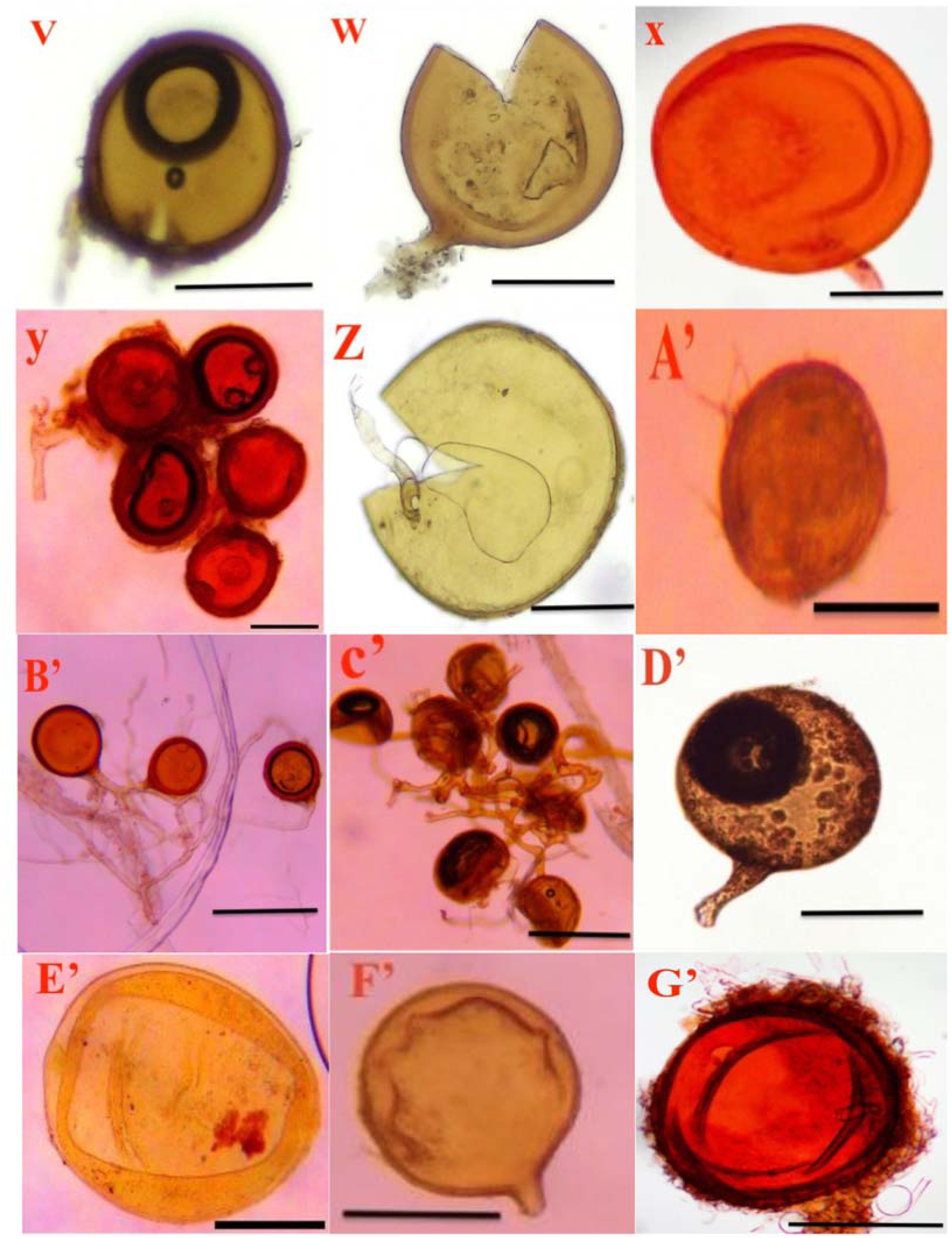
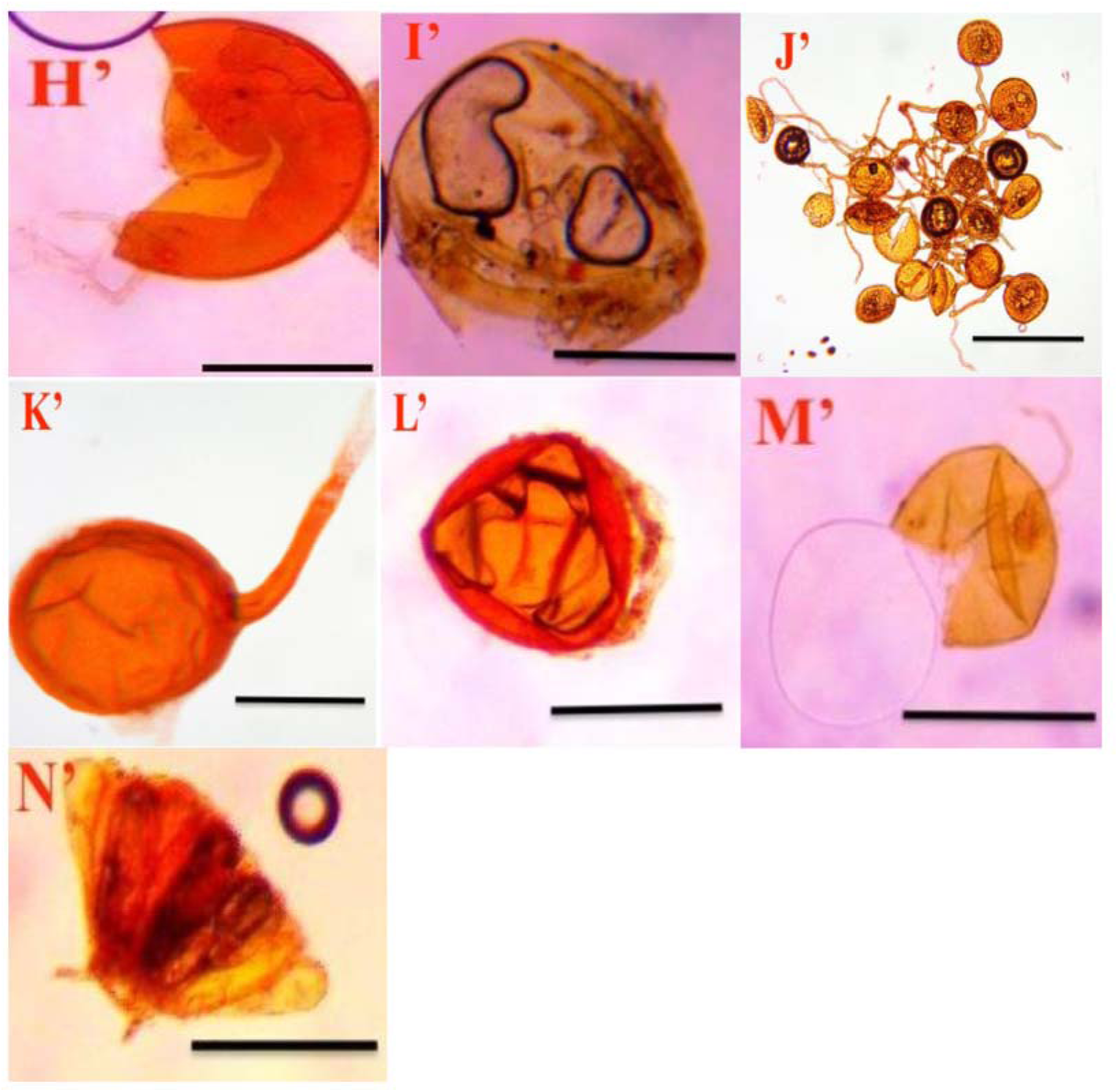
V.) *Glomus ambisporum* W.) *Glomus badium*X*.)GlomuscaledoniumY.)Glomus constrictumZ.)Glomus hoiiA’.)Glomus macrocarpumB’.)Glomus monosporumC’)Glomus velum D’.)Glomus glomerulatumE’.)Glomus occultum F’.) Glomus sps 1G’.)Glomus* sp. 2 *H’.) Glomus etunicatum I’.)Pacispora boliviana J’.)Rhizophagus aggregatus K’.) Rhizophagus irregularis L’.)Scutellospora heterogama M’.)Scutellospora calospora N’.) Sclerocystis sinuosum.* (Scale 40 µm).

**Figure 3.**
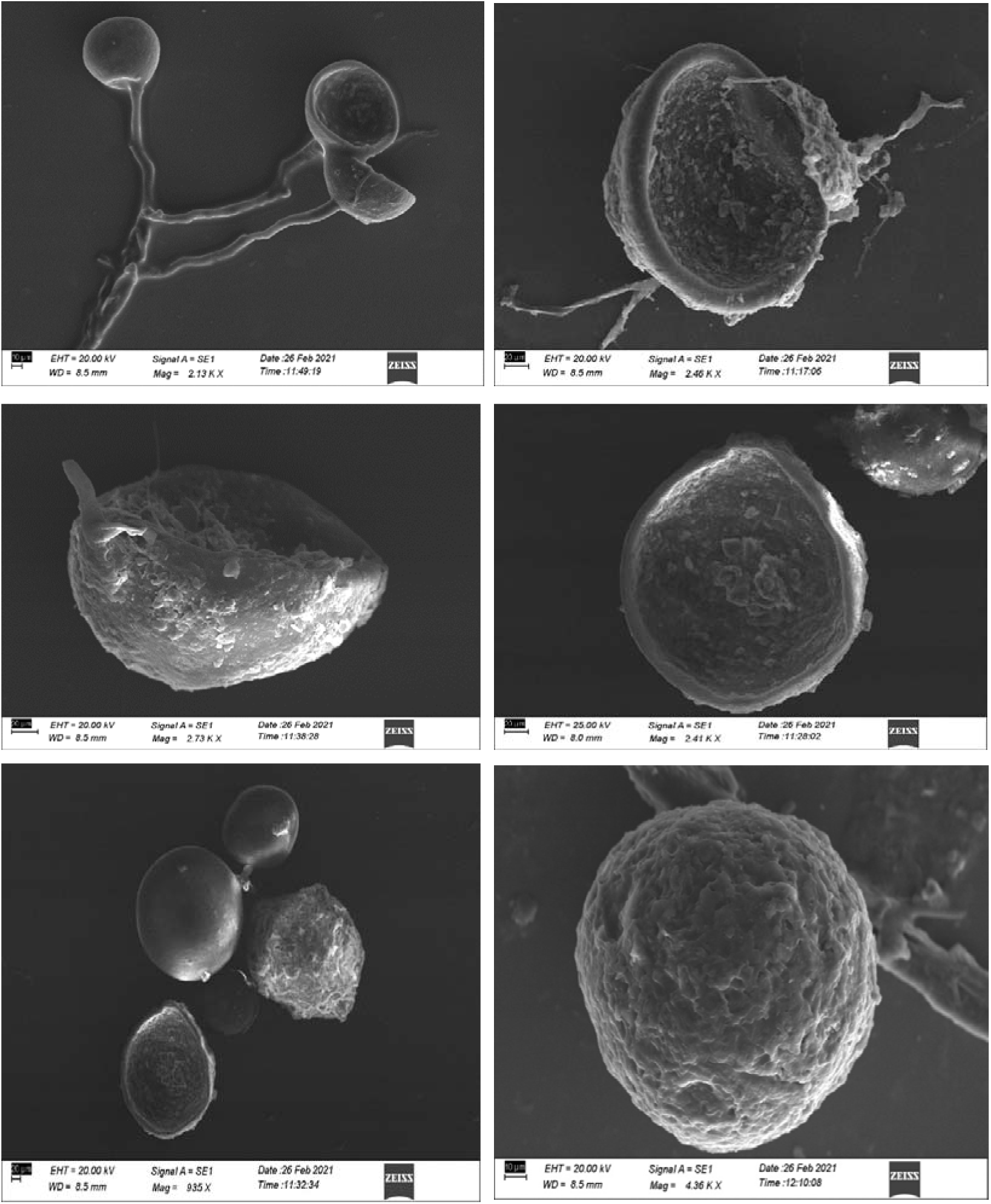
Scanning electron microscopy images of some of the isolated AMF species.

**Figure 4.**
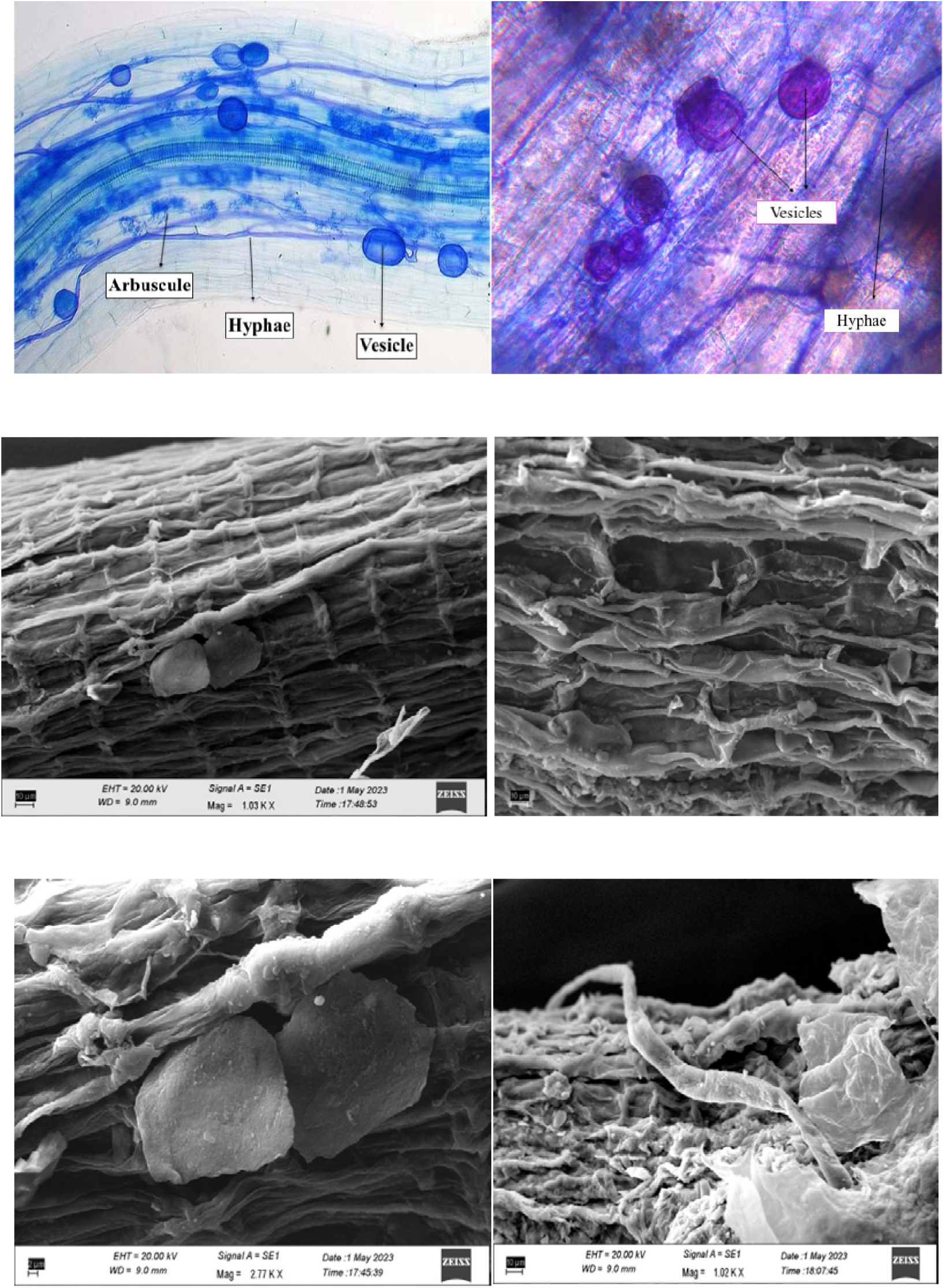
Root colonization images of AMF. In light microscopy and SEM images of arbuscules, vesicles and hyphae are clearly visible.

**Figure 5.**
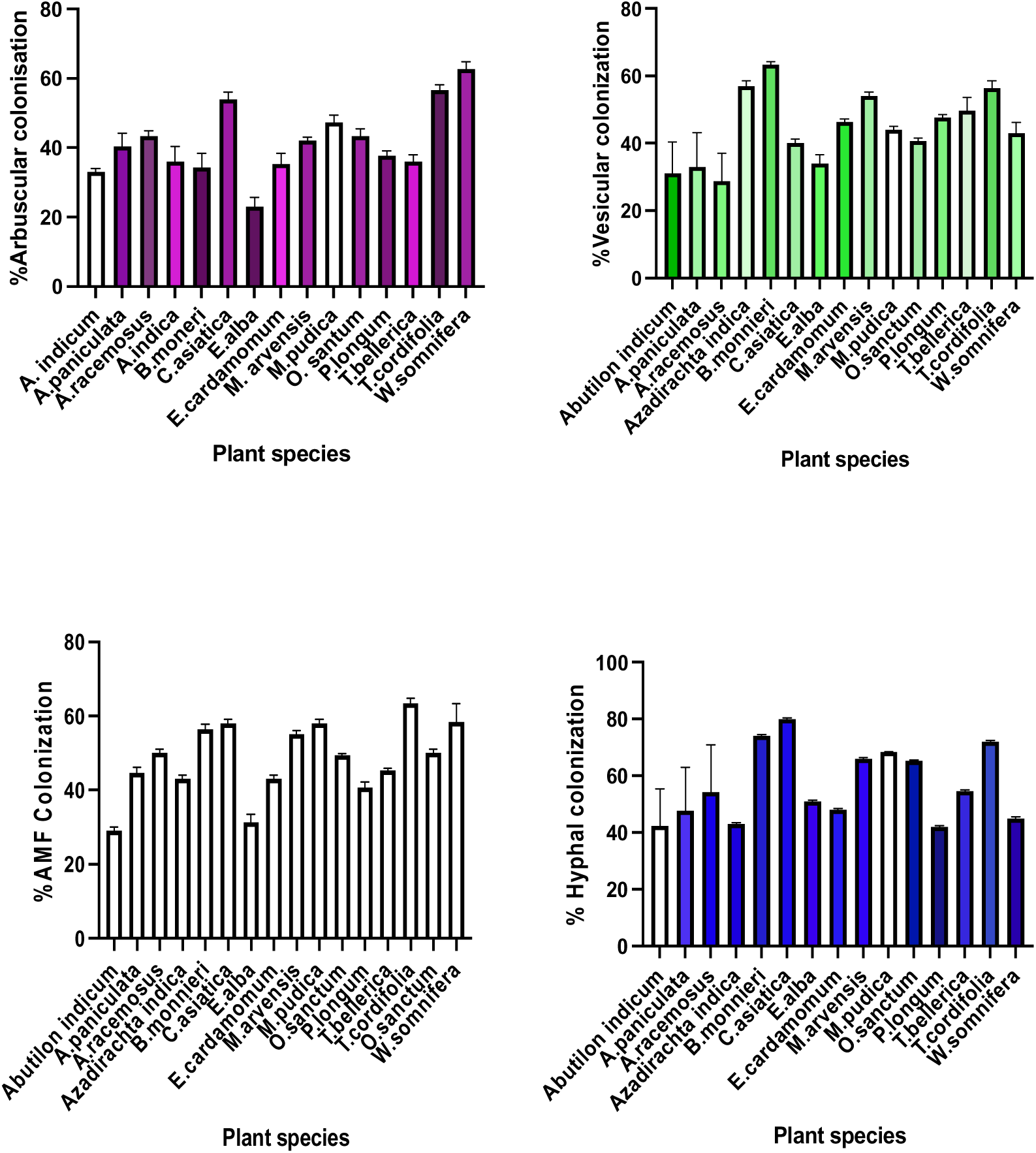
Arbuscular, vesicular, hyphal, and total AMF colonization found in the rhizospheric soil of medicinal plants investigated.

**Figure 6:**
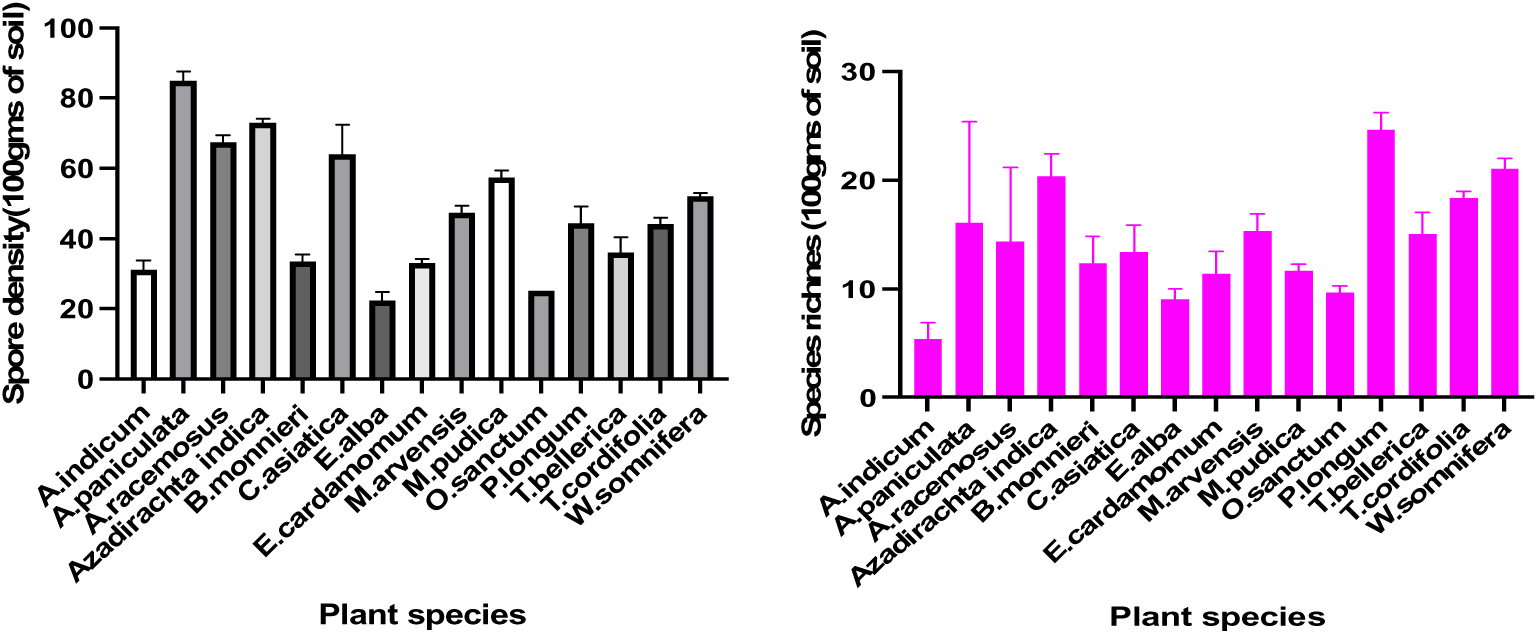
Spore density and species richness of different AMF species isolated from the plants taken for this study.

**Figure 7.**
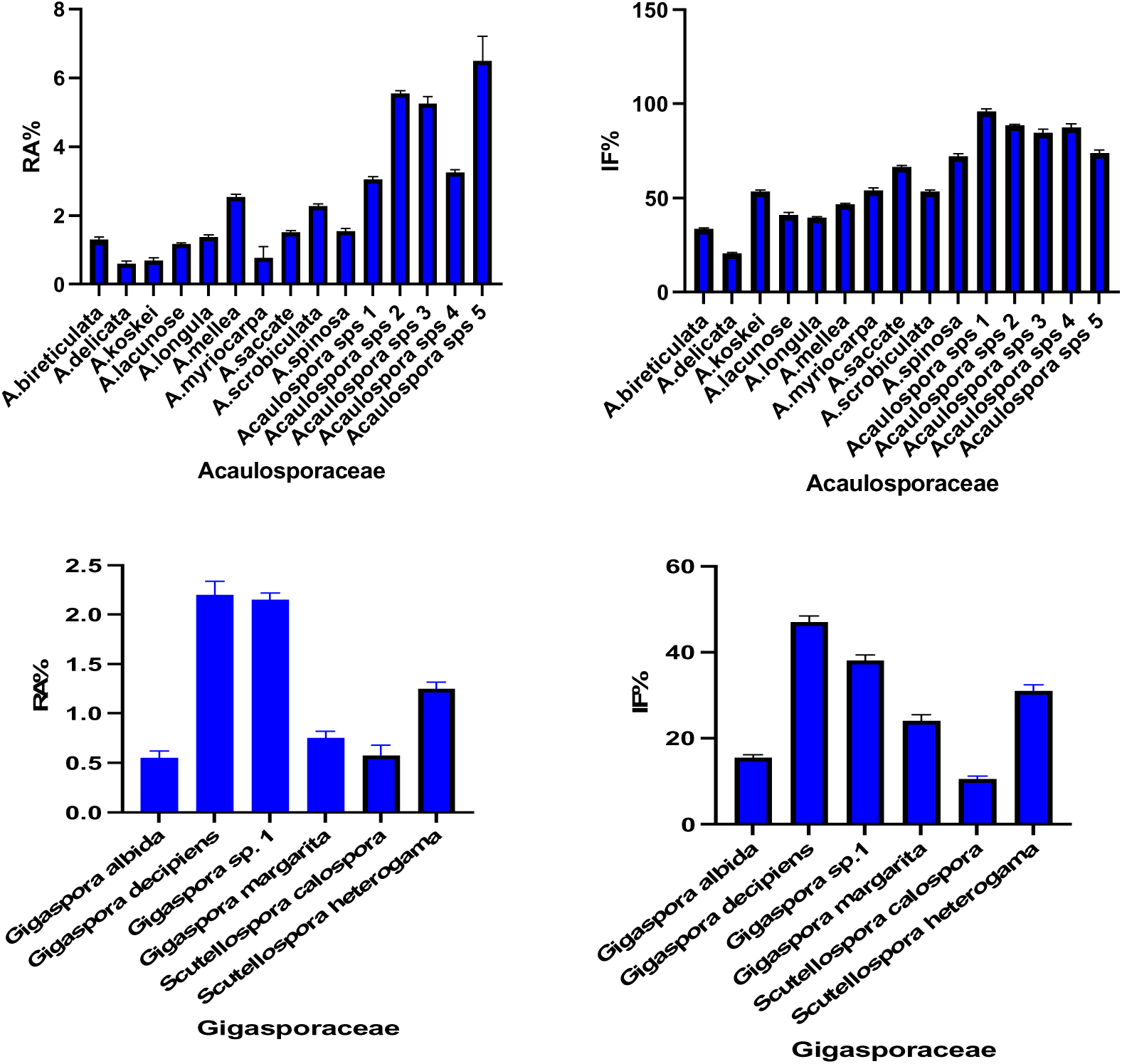
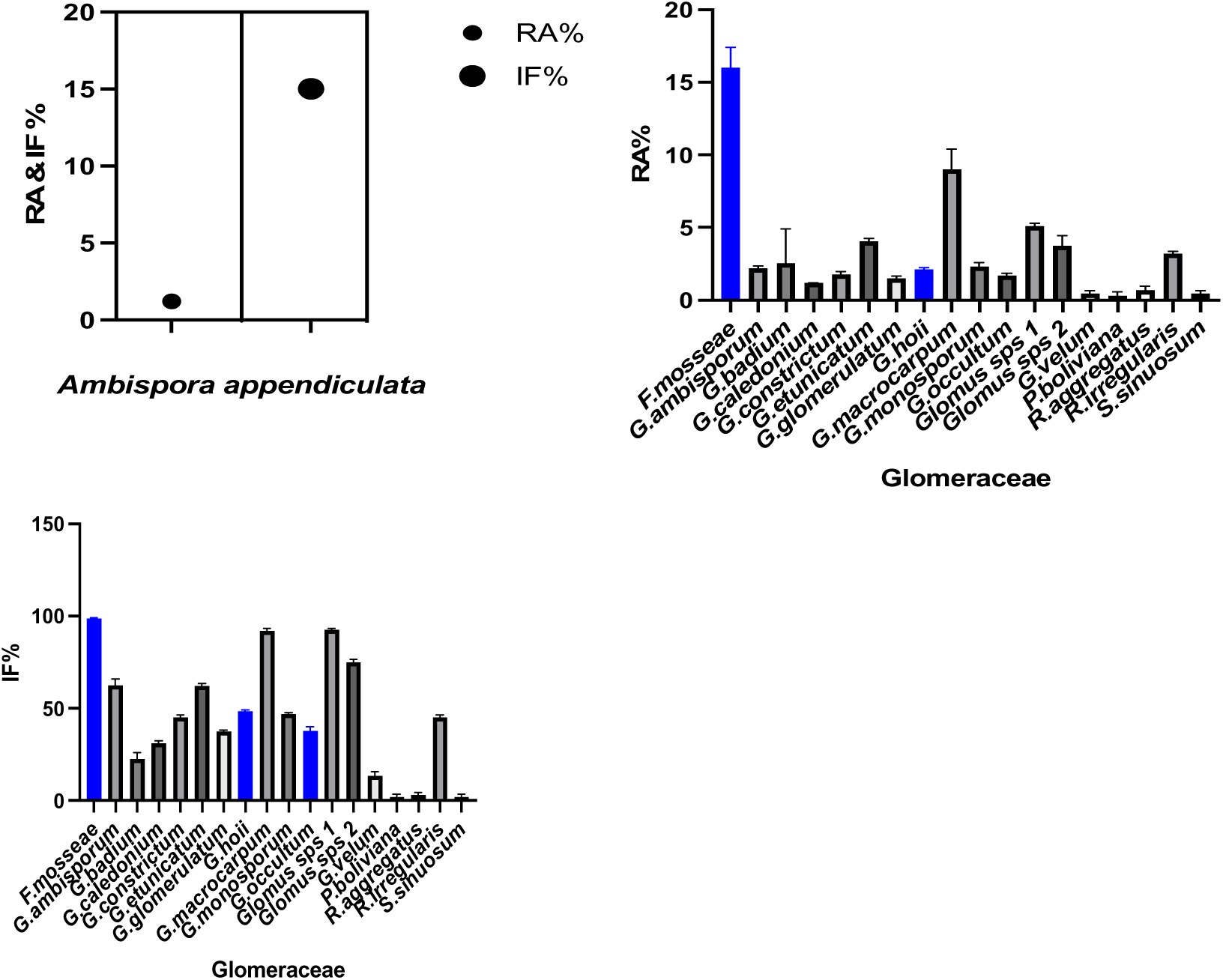
Relative abundance and Isolation frequency of AMF species of Acaulosporaceae, Gigasporaceae, Ambisporaceae and Glomeraceae isolated from rhizospheric soil of medicinal plants investigated.

**Figure 8.**
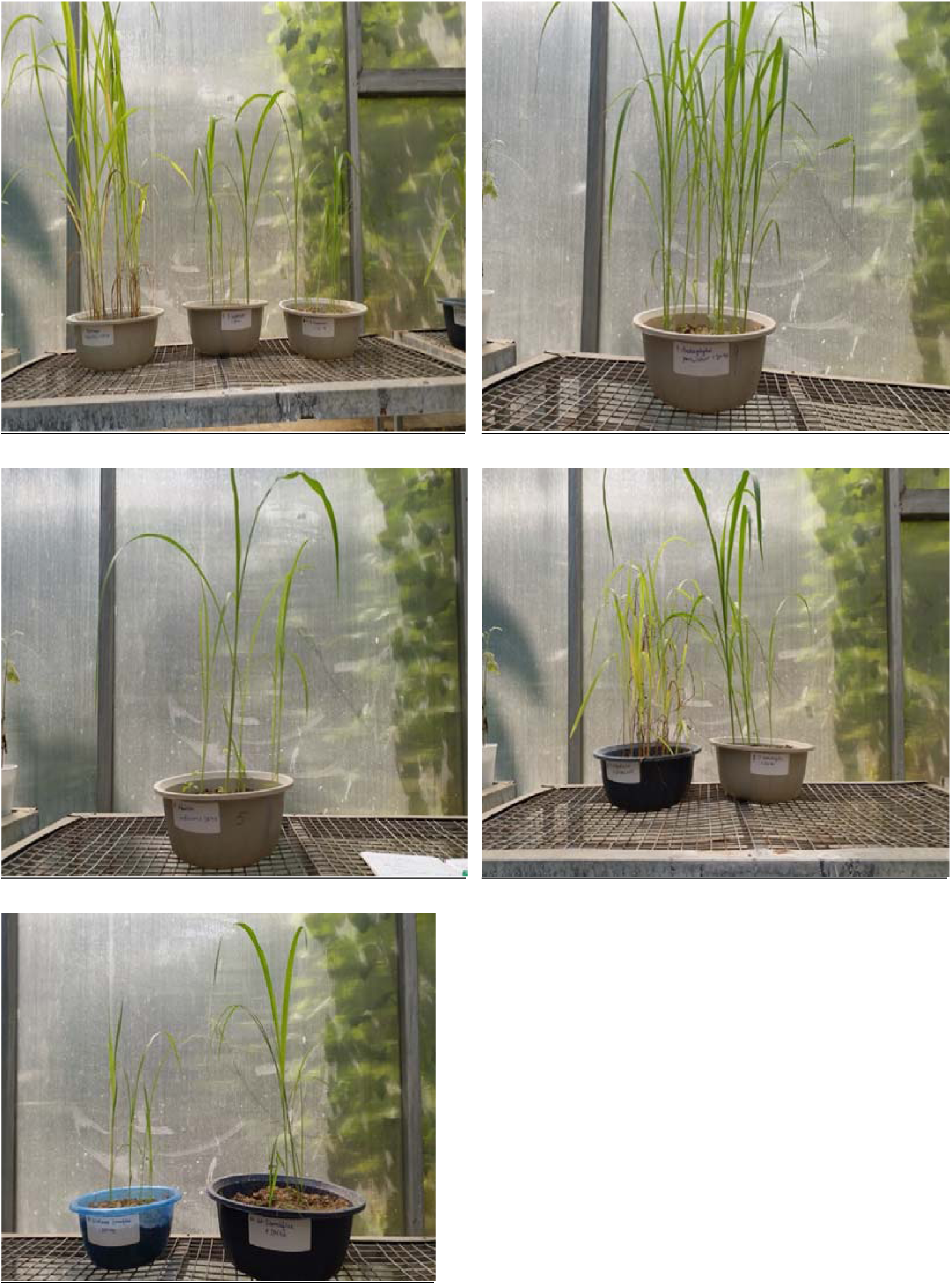
Trap culture images of the rhizospheric soil of some of the selected medicinal plants using *Sorghum bicolor* (SPV40) as host.

### Calculation and statistical analysis

For the purpose of this study various statistical parameters were used such as relative abundance, isolation frequency, Shannon-Wiener index of diversity, Simpson index of dominance and Evenness (Dandan and Zhiwei, 2007). Spore density and species richness were expressed as the total number of spores and the number of species discovered in 100g of soil. Pearson’s correlation coefficient was used to examine the connections between AMF colonization, spore density, and soil physico-chemical parameters. One-way ANOVA was used for statistical analysis, and standard errors of means were determined.

### Scanning electron microscopy

Preparing root samples for visualization of AMF colonization through scanning electron microscopy (SEM) involves several steps to ensure that the samples were properly preserved, dehydrated, and coated for imaging. Following steps were followed for the procedure: -i.) Fixation: Root specimen was cut into small pieces (1-2 mm). The samples were fixed in 2.5% glutaraldehyde in 0.1 M phosphate buffer for 2-4 hours at room temperature or overnight at 4°C. ii.) Wash with Buffer: The samples were rinsed three times with 1 M sodium cacodylate buffer (pH 7.2-7.4) for 15 minutes each to remove excess fixative. iii.)Post-Fixation: The samples were post-fixed in 1% osmium tetroxide in 1 M sodium cacodylate buffer for 1-2 hours at room temperature. This step enhances contrast for SEM imaging. The samples were then washed three times with distilled water. iv.) Dehydration: The samples were then dehydrated through a graded ethanol series (30%, 50%, 70%, 90%, and 100%). Each step was allowed to proceed for 15-30 minutes. A critical point drying apparatus was used to remove the ethanol. This replaced the ethanol with liquid COLJ, preventing structural damage during drying. v.) Mounting: The samples were then mounted on SEM stubs using a conductive adhesive (colloidal silver paint was commonly used) The adhesive was allowed to dry completely. vi.) Coating: The samples were coated with a thin layer of conductive material (gold) using a sputter coater. This step enhances conductivity and reduces charging during SEM imaging. vii.) SEM Imaging: The sample stubs were transferred to the SEM chamber. The imaging parameters (e.g., voltage, working distance) were adjusted based on the instrument specifications. The SEM images of the fungal structures were captured at various magnifications.

**Table 1.**
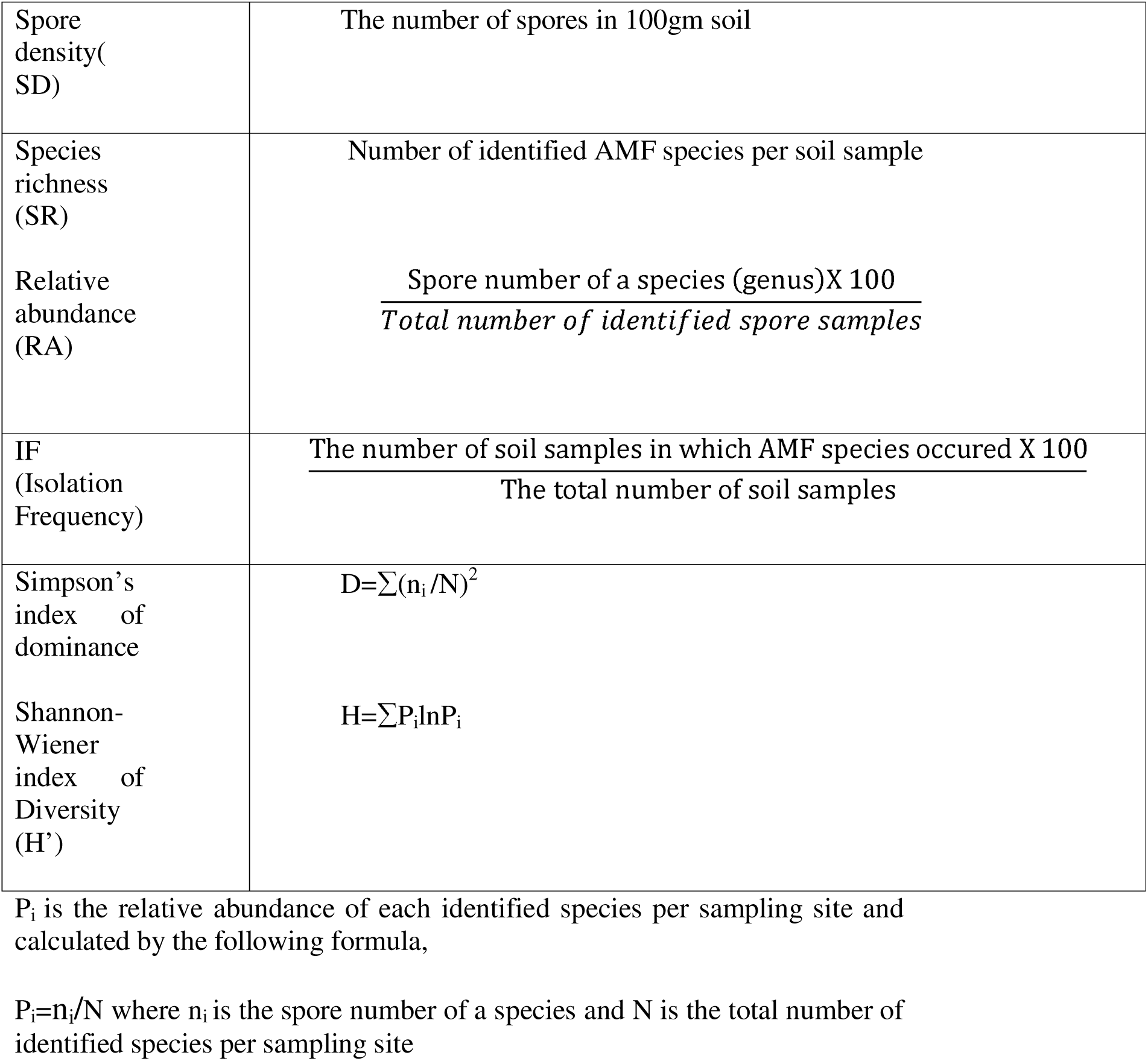
Diversity indices used to describe AMF communities.

Shannon–Wiener index of AMF diversity from sample collected from rhizospheric soil ranged from 1.36 to 2.12 while in trap culture soil sample, it ranged from 2.6 to 3.17. Simpson’s dominance index of AMF ranged from 0.19 to 0.20 from rhizospheric soil, while it ranged from 0.21 to 0.24 in trap culture soil. The value of evenness of AMF species ranged from 0.93 to 0.99 in rhizospheric soil sample while it ranged from 0.81 to 0.91 in trap culture soil.

## Results & Discussion

### AMF colonization

AMF structures, i.e., arbuscules, vesicles and hyphae, and occasionally, intraradical spores were present in the root. The rate of AMF colonization was found to be highest in *Tinospora cordifolia* among the studied plant species. Total AMF colonization ranged from 29% to 63.3% among the studied plants. Arbuscular colonization was found to be highest in *Withania somnifera* followed by *Tinospora cordifolia* and *Centella asiatica.* Vesicular colonization was found to be highest in *Bacopa monnieri* followed by *Tinospora cordifolia* and *Azadirachta indica*. Hyphal colonization was found to be highest *Centella asiatica* followed by *Bacopa monnieri* and *Tinospora cordifolia*. AMF colonization was found to be highest in *Tinospora cordifolia* followed by *Centella asiatica* and *Bacopa monnieri*. Spore density was found to be highest in *Andrographis paniculata* followed by *Azadirachta indica*. Species richness was found to be highest in *Piper longum* followed by *Withania somnifera*. AMF colonization had a significant positive correlation with both moisture content (r = 0.923, p < 0.01), and pH (r = 0.828, p < 0.03). Spore density also showed positive correlation with moisture content (r = 0.933, p < 0.01) and pH (r = 0.958, p < 0.01).

### AMF spore density (SD) and species richness (SR)

AMF species richness and spore density from the rhizospheric soil is presented in figure 6. SD was found to be lowest in *Eclipta alba* (20 spores per 100gm of rhizospheric soil) and highest in *Andrographis paniculata* (84 spores per 100gms of rhizospherc soil). SR was lowest in *Abutilon indicum* and highest in *Piper longum*. In case of trap culture soil, SD was found to be lowest in *Abutilon indicum* (54 spores per 100gm of soil) and highest in *Azadirachta indica* (111 spores per 100gm of soil) and SR was found to be lowest in *Abutilon indicum* and highest in *Withania somnifera* (Figure 9). Spore density was found to be highest for trap culture soil of *Azadirachta indica* followed by *Centella asiatica*. Species richness was found to be highest in trap culture soil of *Withania somnifera* followed by *Mentha arvensis*. Altogether 41 AMF species were isolated from both the rhizospheric and trap culture soil of selected plants (18 belonging to Glomeraceae, 15 belonging to Acaulosporaceae, 6 belonging to Gigasporaceae and 1 to Ambisporaceae). AMF species isolated from the rhizospheric soil of selected plant from both rhizospheric and trap culture derived inoculum with their relative abundance and frequency is presented in figure 6 & 8.

**Figure 9.**
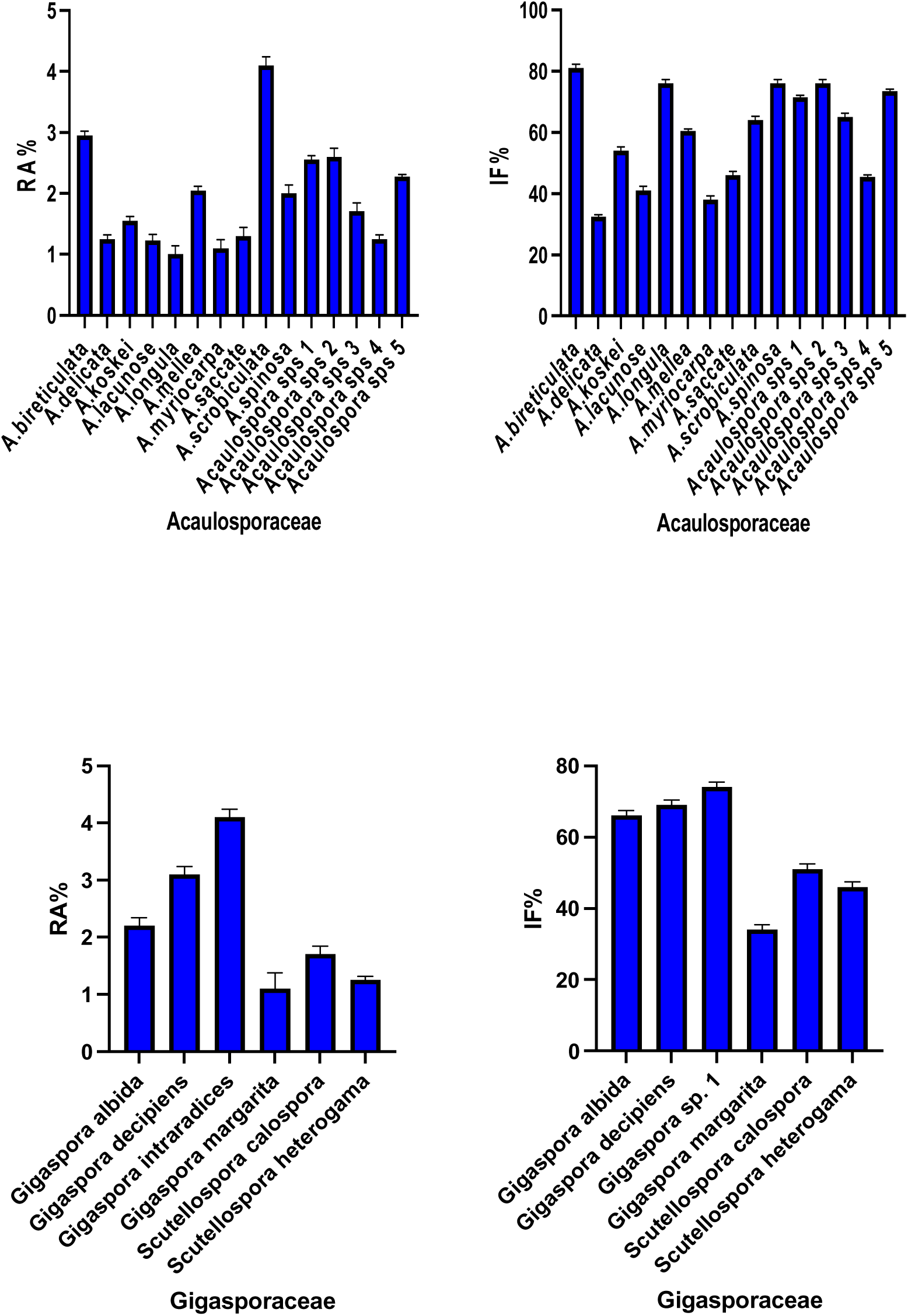
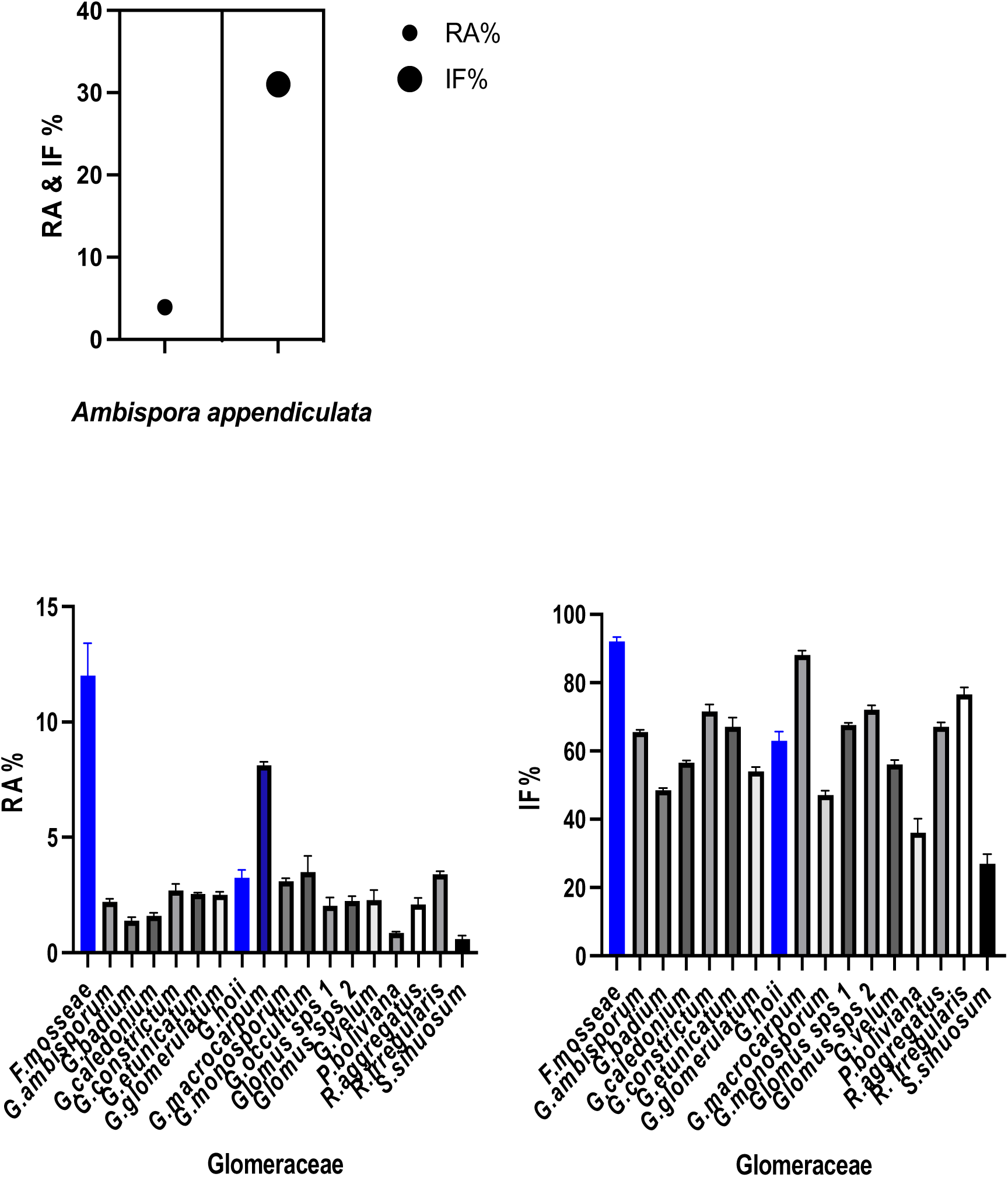
Relative abundance and Isolation frequency of AMF species of Acaulosporaceae, Gigasporaceae, Ambisporaceae and Glomeraceae isolated from trap culture soil.

**Figure 10.**
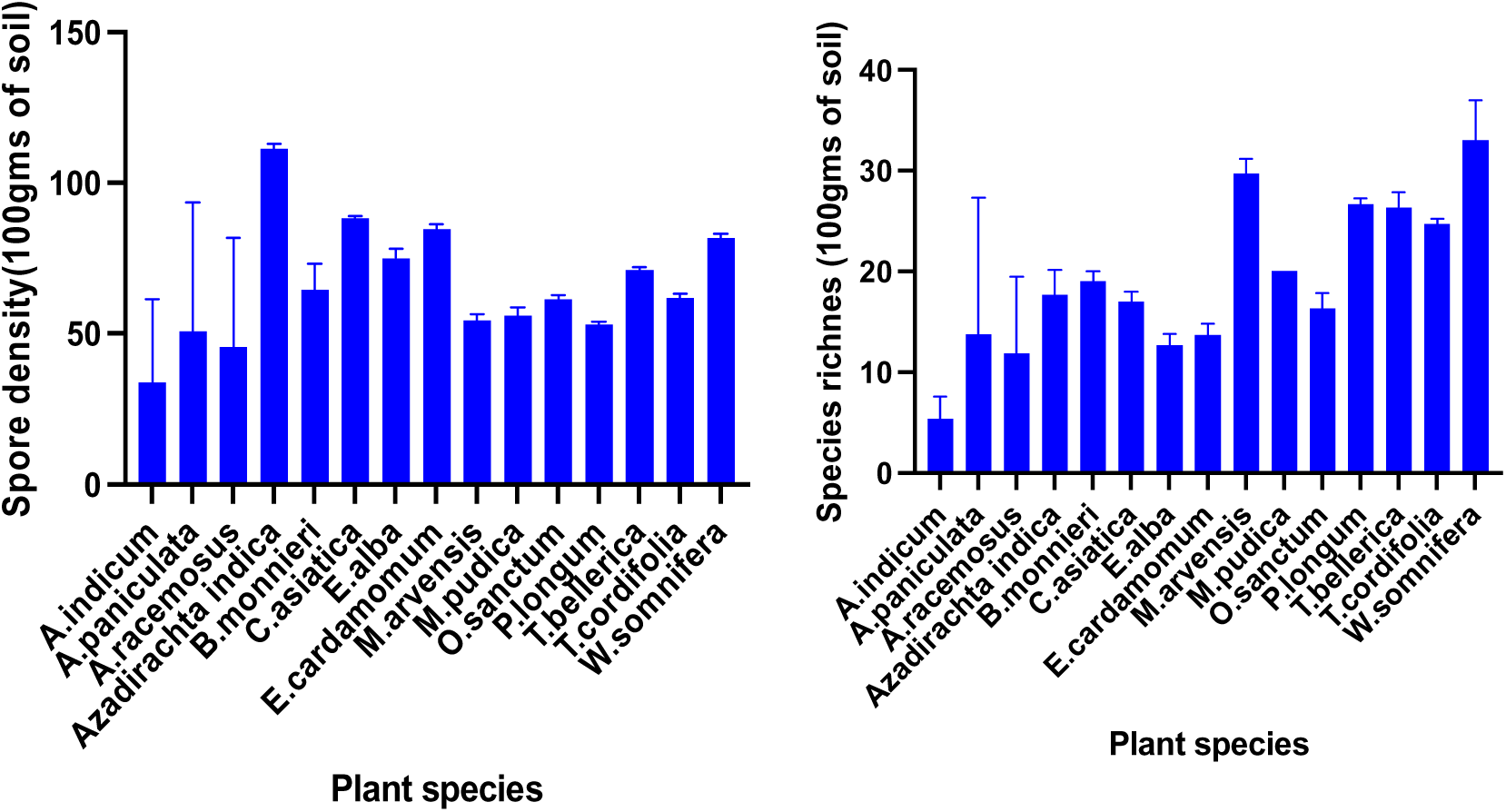
Spore density and species richness of different AMF species isolated from trap culture.

Among Acaulosporaceae the relative abundance (RA%) and isolation frequency (IF%) was found to be highest for *Acaulospora* sp. 5. Among the species of Gigasporaceae the RA% was found to be highest for *Gigaspora intraradices* and IF% was found to be highest for *Gigaspora decipiens*. *Ambispora appendiculata* was the only species of Ambisporaceaea that was found in the studies.

The RA% for *F.mosseae* and *G.macrocarpum* was found to be highest in Glomeraceae. The IF% among Glomeraceae was found to be highest for *F.mosseae*, *G.macrocarpum* and *Glomus* sp. 1.

In trap culture soil the RA% for Acaulosporaceae was found to be highest in *A.scrobiculata* and IF% was found to be highest in *A. bireticulata.* Among Gigasporaceae the RA% and IF% was found to be highest for *Gigaspora* sp.1. Among trap culture soil also *Ambispora appendiculata* was the only species of Ambisporaceae that was found. Among Glomeraceae, RA% and IF% was found to be highest for *F.mosseae* followed by *G.macrocarpum*. Sporulation and distribution of AMF were reflected as a whole according to significant positive correlation between relative abundance and isolation frequency of AMF which in case of Acaulosporaceae (r = 0.94, P<0.0001), Glomeraceae (r=0.8210, P<0.0001) and Gigasporaceae (r=0.9337, P<0.0063) was significant. However, not every dominant species had both high isolation frequency and relative abundance at the same time. *A.mellea* and *G.ambisporum* were found to be widely distributed but had low relative abundance. There was no significant positive correlation between spore density and species richness (r=0.25, P=0.05). Significant positive correlation was also found between relative abundance and isolation frequency in trap culture experiments.

### Soil analysis results

Soil physiochemical properties of botanical garden soil, BHU, Varanasi, district of Eastern Uttar Pradesh was evaluated analysing six rhizosphere soil samples each from every plant studied. In general, soils were alkaline in pH, low in soluble salts and organic carbon. The pH of the rhizospheric soil ranged from 6.9 to 8.5 with a mean of 9.1 indicating neutral to alkaline nature of soil. The organic carbon content ranged 1-5g kg ^-1^.

**Table 1.**
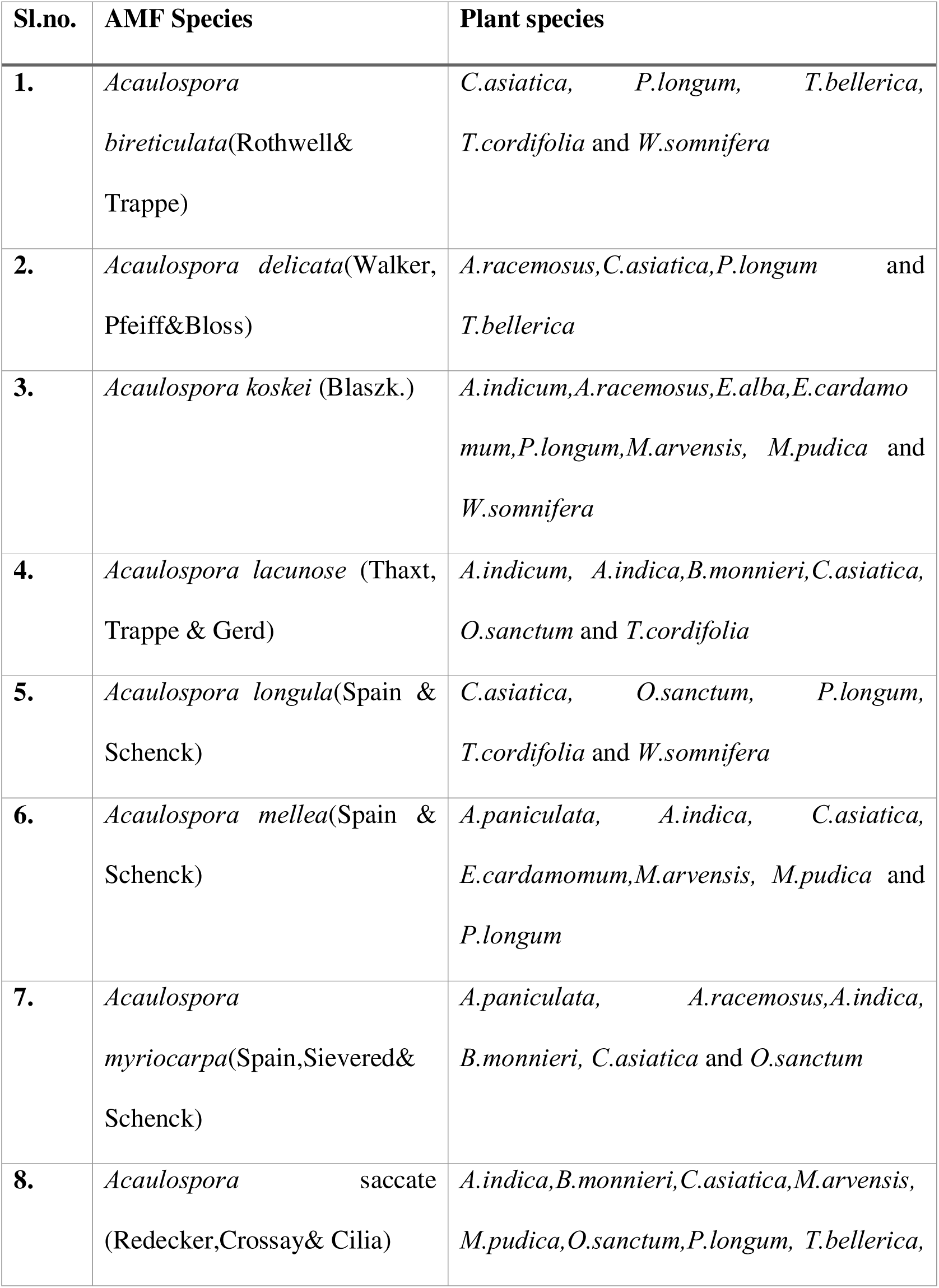
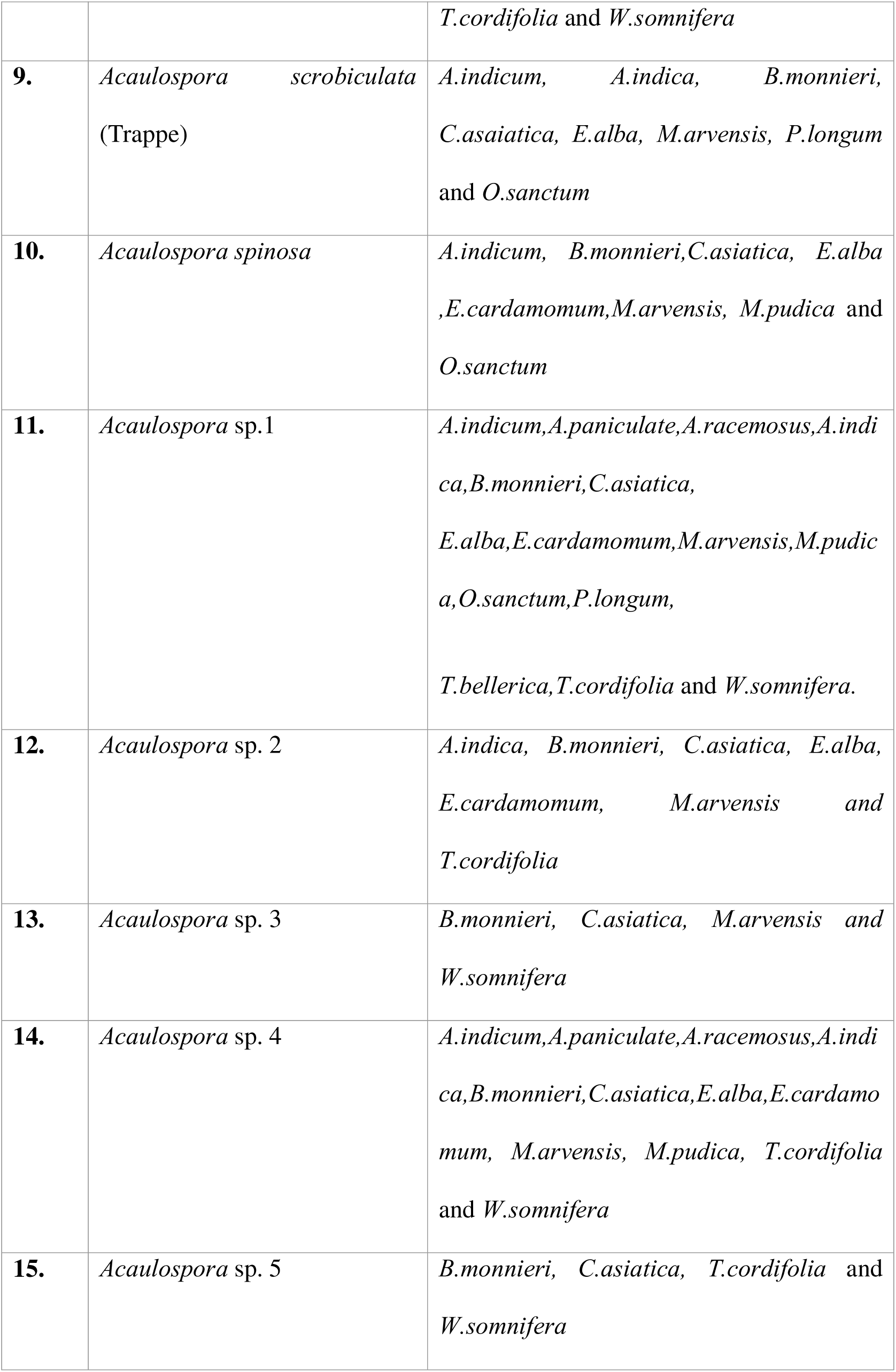
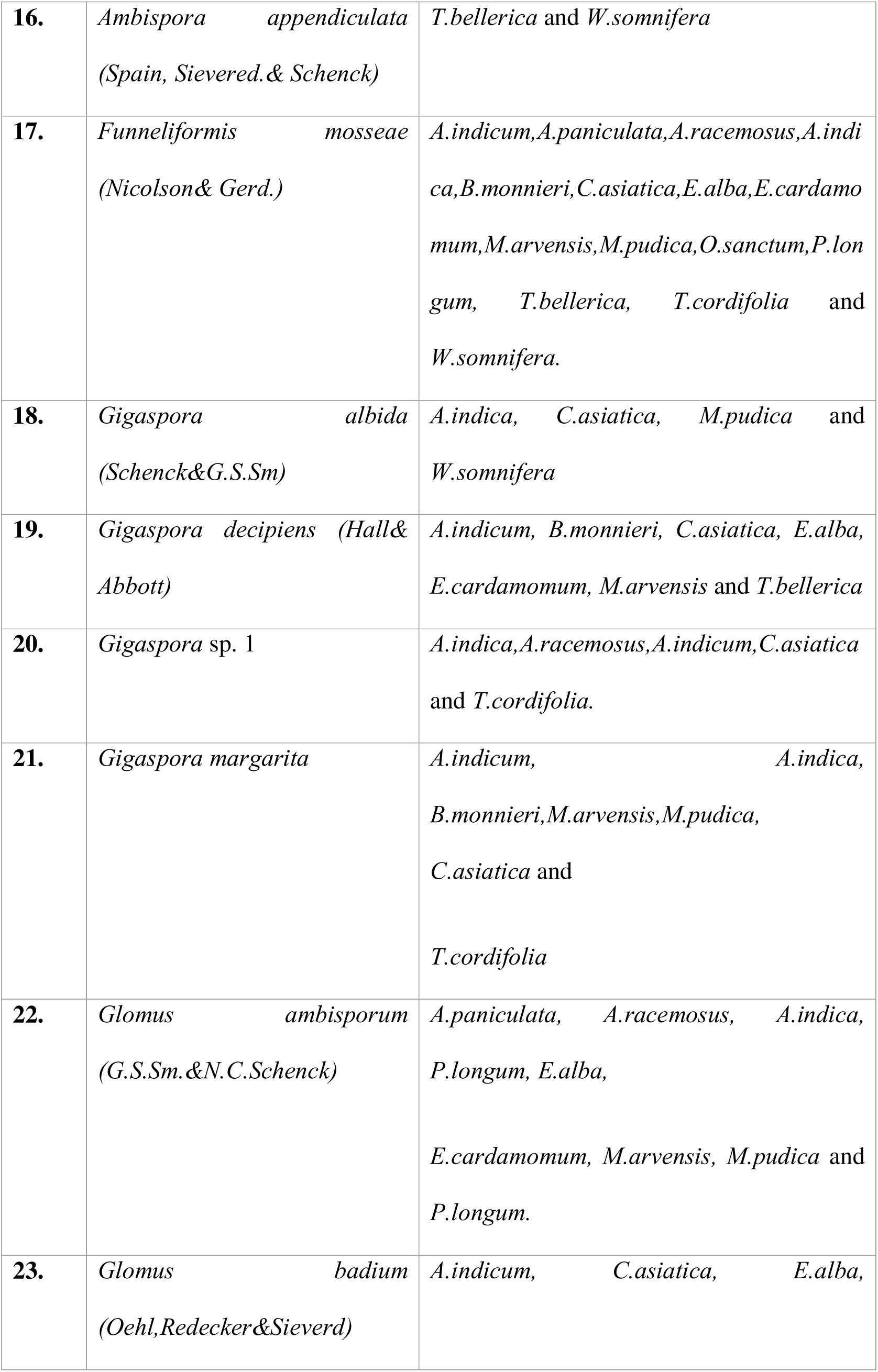
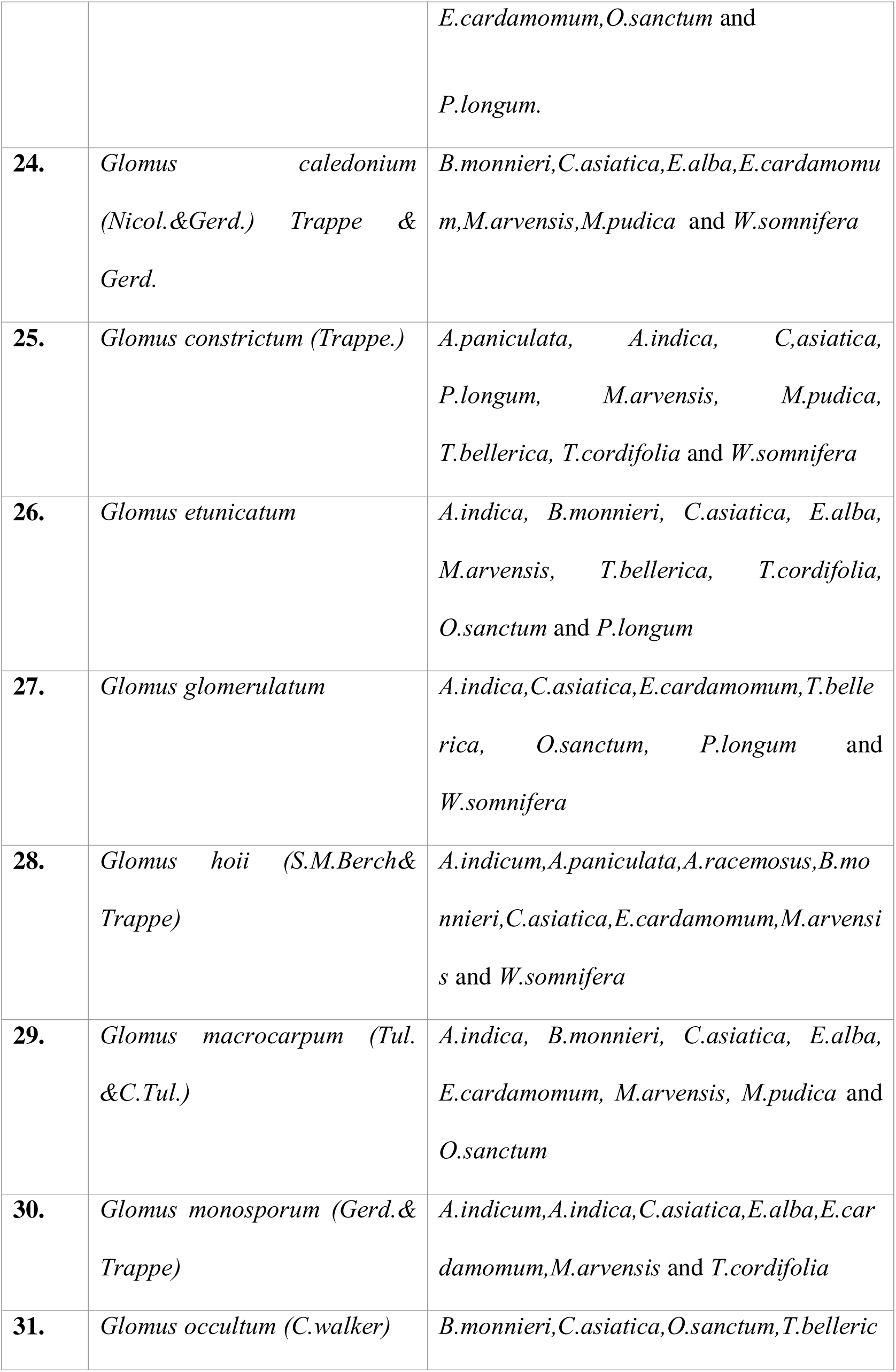
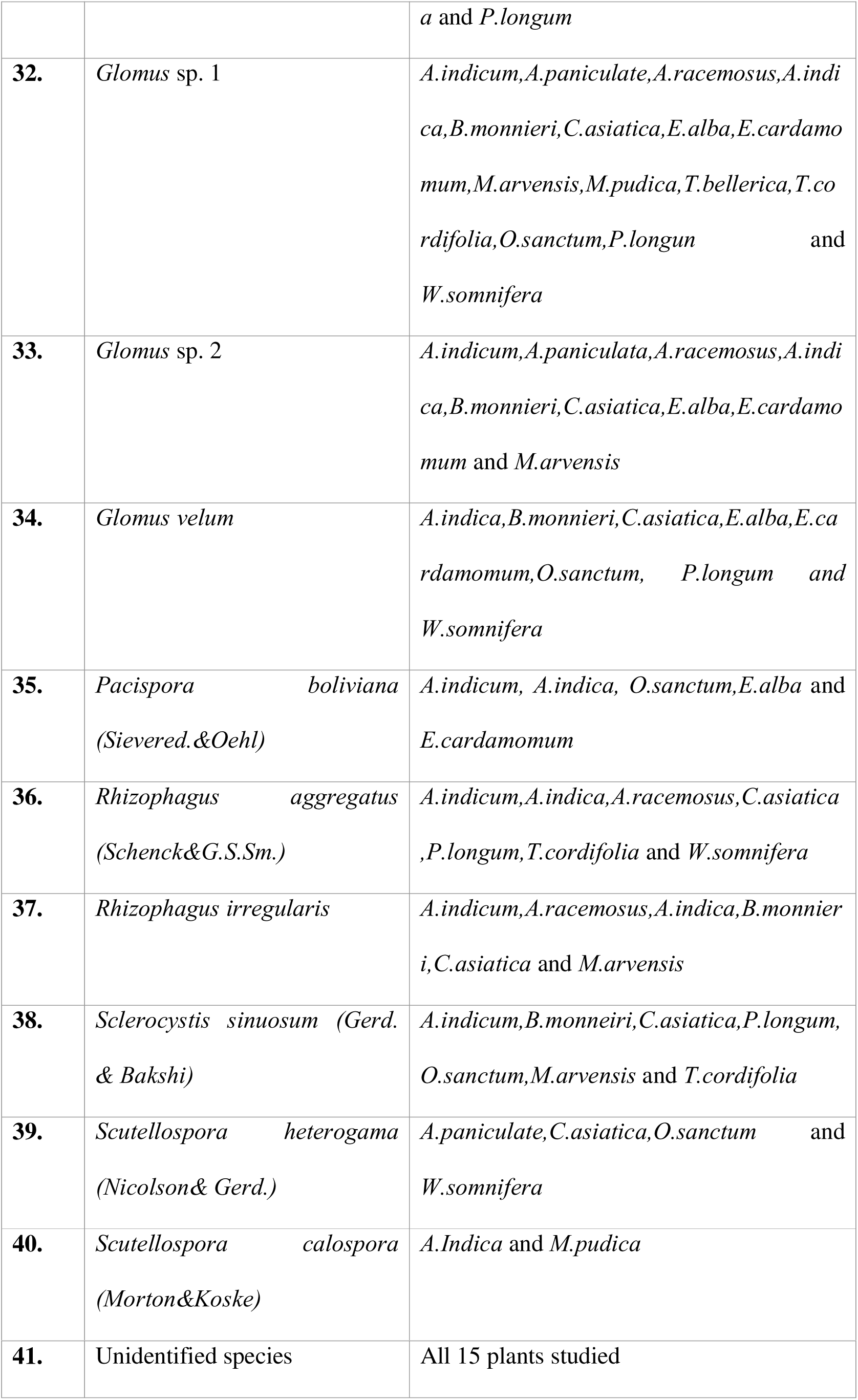
List of isolated AMF species from the rhizosphere of selected medicinal plants.

| Sl.no. | AMF Species | Plant species |
| --- | --- | --- |
| 1. | *Acaulospora*<br>*bireticulata*(Rothwell& Trappe) | *C.asiatica*, *P.longum*, *T.bellerica*,<br>*T.cordifolia* and *W.somnifera* |
| 2. | *Acaulospora delicata*(Walker, Pfeiff&Bloss) | *A.racemosus*,*C.asiatica*,*P.longum* and<br>*T.bellerica* |
| 3. | *Acaulospora koskei* (Blaszk.) | *A.indicum*,*A.racemosus*,*E.alba*,*E.cardamomum*,*P.longum*,*M.arvensis*, *M.pudica* and<br>*W.somnifera* |
| 4. | *Acaulospora lacunose* (Thaxt, Trappe & Gerd) | *A.indicum*, *A.indica*,*B.monneri*,*C.asiatica*,<br>*O.sanctum* and *T.cordifolia* |
| 5. | *Acaulospora longula*(Spain & Schenck) | *C.asiatica*, *O.sanctum*, *P.longum*,<br>*T.cordifolia* and *W.somnifera* |
| 6. | *Acaulospora mellea*(Spain & Schenck) | *A.paniculata*, *A.indica*, *C.asiatica*,<br>*E.cardamomum*,*M.arvensis*, *M.pudica* and<br>*P.longum* |
| 7. | *Acaulospora*<br>*myriocarpa*(Spain,Sievered& Schenck) | *A.paniculata*, *A.racemosus*,*A.indica*,<br>*B.monneri*, *C.asiatica* and *O.sanctum* |
| 8. | *Acaulospora* saccate<br>(Redecker,Crossay& Cilia) | *A.indica*,*B.monneri*,*C.asiatica*,*M.arvensis*,<br>*M.pudica*,*O.sanctum*,*P.longum*, *T.bellerica*, |
|  |  | *T.cordifolia* and *W.somnifera* |
| **9.** | *Acaulospora scrobiculata*<br>(Trappe) | *A.indicum*, *A.indica*, *B.monneri*,<br>*C.asaiatica*, *E.alba*, *M.arvensis*, *P.longum*<br>and *O.sanctum* |
| **10.** | *Acaulospora spinosa* | *A.indicum*, *B.monneri*, *C.asiatica*, *E.alba*<br>, *E.cardamomum*, *M.arvensis*, *M.pudica* and<br>*O.sanctum* |
| **11.** | *Acaulospora* sp.1 | *A.indicum*, *A.paniculate*, *A.racemosus*, *A.indi*<br>*ca*, *B.monneri*, *C.asiatica*,<br>*E.alba*, *E.cardamomum*, *M.arvensis*, *M.pudic*<br>*a*, *O.sanctum*, *P.longum*,<br>*T.bellerica*, *T.cordifolia* and *W.somnifera*. |
| **12.** | *Acaulospora* sp. 2 | *A.indica*, *B.monneri*, *C.asiatica*, *E.alba*,<br>*E.cardamomum*, *M.arvensis* and<br>*T.cordifolia* |
| **13.** | *Acaulospora* sp. 3 | *B.monneri*, *C.asiatica*, *M.arvensis* and<br>*W.somnifera* |
| **14.** | *Acaulospora* sp. 4 | *A.indicum*, *A.paniculate*, *A.racemosus*, *A.indi*<br>*ca*, *B.monneri*, *C.asiatica*, *E.alba*, *E.cardamo*<br>*mum*, *M.arvensis*, *M.pudica*, *T.cordifolia*<br>and *W.somnifera* |
| **15.** | *Acaulospora* sp. 5 | *B.monneri*, *C.asiatica*, *T.cordifolia* and<br>*W.somnifera* |
| **16.** | *Ambispora appendiculata*<br>(Spain, Sievered. & Schenck) | *T.bellerica* and *W.somnifera* |
| **17.** | *Funneliformis mosseae*<br>(Nicolson & Gerd.) | *A.indicum*, *A.paniculata*, *A.racemosus*, *A.indica*, *B.monneri*, *C.asiatica*, *E.alba*, *E.cardamomum*, *M.arvensis*, *M.pudica*, *O.sanctum*, *P.longum*, *T.bellerica*, *T.cordifolia* and *W.somnifera*. |
| **18.** | *Gigaspora albida*<br>(Schenck & G.S.Sm) | *A.indica*, *C.asiatica*, *M.pudica* and *W.somnifera* |
| **19.** | *Gigaspora decipiens* (Hall & Abbott) | *A.indicum*, *B.monneri*, *C.asiatica*, *E.alba*, *E.cardamomum*, *M.arvensis* and *T.bellerica* |
| **20.** | *Gigaspora* sp. 1 | *A.indica*, *A.racemosus*, *A.indicum*, *C.asiatica* and *T.cordifolia*. |
| **21.** | *Gigaspora margarita* | *A.indicum*, *A.indica*, *B.monneri*, *M.arvensis*, *M.pudica*, *C.asiatica* and *T.cordifolia* |
| **22.** | *Glomus ambisporum*<br>(G.S.Sm. & N.C.Schenck) | *A.paniculata*, *A.racemosus*, *A.indica*, *P.longum*, *E.alba*, *E.cardamomum*, *M.arvensis*, *M.pudica* and *P.longum*. |
| **23.** | *Glomus badium*<br>(Oehl, Redecker & Sieverd) | *A.indicum*, *C.asiatica*, *E.alba*, |
|  |  | *E.cardamomum*, *O.sanctum* and<br>*P.longum*. |
| **24.** | *Glomus caledonium*<br>(Nicol. & Gerd.) Trappe &<br>Gerd. | *B.monneri*, *C.asiatica*, *E.alba*, *E.cardamomu*<br>*m*, *M.arvensis*, *M.pudica* and *W.somnifera* |
| **25.** | *Glomus constrictum* (Trappe.) | *A.paniculata*, *A.indica*, *C.asiatica*,<br>*P.longum*, *M.arvensis*, *M.pudica*,<br>*T.bellerica*, *T.cordifolia* and *W.somnifera* |
| **26.** | *Glomus etunicatum* | *A.indica*, *B.monneri*, *C.asiatica*, *E.alba*,<br>*M.arvensis*, *T.bellerica*, *T.cordifolia*,<br>*O.sanctum* and *P.longum* |
| **27.** | *Glomus glomerulatum* | *A.indica*, *C.asiatica*, *E.cardamomum*, *T.belle*<br>*rica*, *O.sanctum*, *P.longum* and<br>*W.somnifera* |
| **28.** | *Glomus hoii* (S.M.Berch &<br>Trappe) | *A.indicum*, *A.paniculata*, *A.racemosus*, *B.mo*<br>*nnieri*, *C.asiatica*, *E.cardamomum*, *M.arvensi*<br>*s* and *W.somnifera* |
| **29.** | *Glomus macrocarpum* (Tul.<br>& C.Tul.) | *A.indica*, *B.monneri*, *C.asiatica*, *E.alba*,<br>*E.cardamomum*, *M.arvensis*, *M.pudica* and<br>*O.sanctum* |
| **30.** | *Glomus monosporum* (Gerd. &<br>Trappe) | *A.indicum*, *A.indica*, *C.asiatica*, *E.alba*, *E.car*<br>*damomum*, *M.arvensis* and *T.cordifolia* |
| **31.** | *Glomus occultum* (C.walker) | *B.monneri*, *C.asiatica*, *O.sanctum*, *T.belleric* |
|  |  | *a* and *P.longum* |
| 32. | *Glomus* sp. 1 | *A.indicum,A.paniculate,A.racemosus,A.indica,B.monneri,C.asiatica,E.alba,E.cardamomum,M.arvensis,M.pudica,T.bellerica,T.cordifolia,O.sanctum,P.longum* and *W.somnifera* |
| 33. | *Glomus* sp. 2 | *A.indicum,A.paniculata,A.racemosus,A.indica,B.monneri,C.asiatica,E.alba,E.cardamomum* and *M.arvensis* |
| 34. | *Glomus velum* | *A.indica,B.monneri,C.asiatica,E.alba,E.cardamomum,O.sanctum, P.longum* and *W.somnifera* |
| 35. | *Pacispora boliviana* (Sievered.&Oehl) | *A.indicum, A.indica, O.sanctum,E.alba* and *E.cardamomum* |
| 36. | *Rhizophagus aggregatus* (Schenck&G.S.Sm.) | *A.indicum,A.indica,A.racemosus,C.asiatica ,P.longum,T.cordifolia* and *W.somnifera* |
| 37. | *Rhizophagus irregularis* | *A.indicum,A.racemosus,A.indica,B.monneri,C.asiatica* and *M.arvensis* |
| 38. | *Sclerocystis sinuosum* (Gerd. & Bakshi) | *A.indicum,B.monneri,C.asiatica,P.longum, O.sanctum,M.arvensis* and *T.cordifolia* |
| 39. | *Scutellospora heterogama* (Nicolson& Gerd.) | *A.paniculate,C.asiatica,O.sanctum* and *W.somnifera* |
| 40. | *Scutellospora calospora* (Morton&Koske) | *A.Indica* and *M.pudica* |
| 41. | Unidentified species | All 15 plants studied |

## Discussion

The plants selected for this study finds good applications in varied industries including pharmaceutical, herbal, agricultural, food as well as cosmetic industries. These plants have ethnobotanical values and been in use by people of banaras region since time immemorial. Research on AMF has mostly revolved around the identification of AMF associations in different plant communities and their distribution patterns across different habitats in India. The result of this study showed that biodiversity of AMF differ from different plants and the extent of AMF colonization is controlled by the host plants as well as environmental conditions. In our study, the diversity and distribution was described with the help of morphological features.

The result showed the dominancy of genus *Acaulospora* followed by *Glomus,* i.e relatively produced more spores than *Funneliformis, Rhizophagus, Scutellospora* species in the same environmental condition. The degree of colonization and spore density varied markedly among plant species. Estimating AMF root colonization provides valuable insights into the extent and effectiveness of the symbiotic relationship between these fungi and plant roots. It enhances our understanding of the role of AMF in nutrient acquisition, plant growth, and ecosystem functioning. Additionally, monitoring changes in root colonization over time or under different environmental conditions can provide insights into the dynamics of this important mutualistic relationship.

